# Preferential CDR masking in paired antibody language models improves binding affinity prediction

**DOI:** 10.1101/2025.10.31.685149

**Authors:** Mahtab Talaei, Kenji C. Walker, Boran Hao, Eliot Jolley, Yeping Jin, Dima Kozakov, John Misasi, Sandor Vajda, Ioannis Ch. Paschalidis, Diane Joseph-McCarthy

## Abstract

**Background:** Therapeutic antibodies are a leading class of biologics, yet their unique architecture poses challenges for computational modeling. Each antibody comprises paired heavy and light variable domains with conserved framework regions that maintain structure and hypervariable complementarity-determining regions (CDRs) that directly contact antigens. This functional asymmetry, where CDRs determine binding specificity while frameworks provide scaffolding, suggests that region-aware training strategies could yield superior representations. Existing protein language models treat all regions uniformly, potentially missing critical features present in CDRs.

**Methods:** We developed a region-aware pretraining strategy for paired variable domain sequences using two protein language models: a 3 billion parameter model (ESM2) and a compact 600 million parameter model (ESM C). We compared three masking approaches: uniform whole-chain masking, CDR-focused masking, and a hybrid strategy. Final models were trained on over 1.6 million paired antibody sequences and evaluated on binding affinity datasets with over 90,000 antibody variants across six antigens, including single-mutant panels and combinatorial libraries.

**Results:** Here we show that CDR-focused training produces embeddings with superior predictive performance for antibody–antigen binding. Our approach achieves up to 27% improvements in binding affinity prediction compared to benchmarked antibody models. Remarkably, training exclusively on paired sequences proves sufficient; pretraining on billions of unpaired sequences provides no measurable benefit. Our compact model matches or exceeds larger antibody-specific baselines.

**Conclusions:** These findings establish that prioritizing paired sequences with CDR-aware supervision over scale and complex training schemes achieves both computational efficiency and predictive accuracy, providing a practical framework for next generation antibody language models.

## 1 Introduction

Therapeutic antibodies represent one of the most successful classes of biologics, with over 100 FDA-approved treatments and hundreds more in clinical development [1]. These Y-shaped immune proteins achieve remarkable specificity through their modular architecture: paired heavy and light chain variable domains (VH and VL) that together create the antigen-binding site. Within each variable domain, three *complementarity-determining regions (CDRs)* form hypervariable loops that directly contact antigens, while four *framework regions (FRs)* maintain the structural integrity that orients the CDRs toward one another forming the antigen-binding site. Antibody residues that interact with the antigen comprise the paratope, while antigen residues that interact with the antibody form the epitope. This elegant division of labor, where highly variable CDRs evolve extreme sequence diversity for recognition while FRs are minimally changed, presents a fundamental challenge for computational modeling: how to effectively capture both the conserved structural scaffold and the hypervariable functional regions within a unified representation framework.

The expanding clinical impact of antibodies, with dozens of annual FDA approvals across oncology, autoimmunity, and infectious diseases [2, 3], has catalyzed the collection of massive sequence databases [4, 5]. The *Observed Antibody Space (OAS)* contains billions of repertoire sequences from immune responses, while structural databases such as SAbDab provide thousands of experimentally resolved antibody–antigen complexes [6, 7]. This data availability, coupled with a sustained interest in developing antibody biologics driven by clinical successes, creates an unprecedented opportunity for machine learning approaches to accelerate antibody design. However, key challenges remain: most repertoire data lack heavy–light pairing information, CDR sequences exhibit extreme diversity that standard protein models struggle to capture, and the relationship between sequence variation and binding affinity remains to be better elucidated despite recent benchmarking efforts [8, 9].

*Protein language models (PLMs)* offer a promising approach to address these challenges by learning rich representations from raw sequences [10]. ESM2 demonstrated that large-scale *masked language modeling (MLM)* yields embeddings predictive of structure and function, and supports structure prediction (ESMFold) [11]. ESM3 extends this paradigm to multimodal generation across sequence, structure, and function [12]. The parallel ESM Cambrian (ESM C) model family scales up data and training compute to focus on creating representations of the underlying biology of proteins, delivering dramatic performance improvements over ESM2 [13]. While these models provide powerful protein priors for downstream design and prediction [14], antibodies violate key assumptions underlying general protein models: their function concentrates in just six CDR loops (~ 20% of residues), their sequences require proper heavy–light chain pairing for activity, and their CDRs exhibit diversity patterns shaped by somatic hypermutation more than evolutionary conservation. Notably, general PLMs such as ESM2 and ESM C are pretrained on single-chain protein sequences that lack paired sequence information, and thus do not natively capture the heavy–light interaction dependencies critical for antibody function. These unique characteristics suggest that antibody-specific training strategies, particularly those that account for the functional asymmetry between CDRs and frameworks, could yield superior representations for therapeutic design.

Specialized antibody language models address parts of this mismatch. AntiB-ERTy organizes repertoires along affinity-maturation trajectories [15]; AbLang corrects sequencing artifacts that confound generic PLMs [16] and AntiBERTa adapts RoBERTa for paratope prediction [17]. More recent large-scale efforts combine billions of unpaired chains with millions of paired VH–VL sequences (e.g., IgBERT, IgT5) [18], and curriculum/mixture schedules have been proposed to interleave unpaired with paired data, aided by rotary positional embeddings (RoPE) and chain separators for more robust cross-chain attention [19]. While these models leverage antibody-specific architectures and sophisticated training curricula, most employ uniform masking that treats all regions equally during pretraining and may not optimally capture the functional determinants of antibody specificity.

Several recent studies have begun to rebalance self-supervised training toward the more functionally relevant residues that uniform masked language modeling underemphasizes. Olsen et al. [20] developed AbLang2 to address the germline frequency bias, where most masked tokens are easy-to-predict germline residues, by using focal loss to reweight learning toward non-germline (mutated) residues while maintaining germline accuracy. Ng et al. [21] modified the masking distribution itself, preferentially masking CDR3 loops to improve modeling of these high-entropy loops and enhance tasks like native pairing discrimination and binding specificity classification. Gao et al. [22] emphasized CDR generation for antigen-specific design rather than systematically characterizing region-specific trade-offs for transferable embeddings. However, it remains unclear whether systematic CDR-focused masking across all CDRs, applied to paired-only training from the outset, can consistently improve binding affinity prediction without computationally intensive unpaired pretraining.

We address this gap through systematic comparison of masking strategies (whole-chain, CDR-focused, hybrid) on paired sequences across two model scales, evaluated on both region-stratified masked recovery and comprehensive binding affinity benchmarks spanning diverse antigens and mutational contexts.

### Contributions

We systematically test region-aware pretraining on paired VH– VL sequences across two architectures with different scales (ESM2-3B and ESM Cambrian-600M). Our approach progresses from whole-chain (WC) masking, where 15% of all residues are randomly masked during training, to CDR-focused training, where 50% of CDR residues are randomly masked, with a hybrid variant that retains framework (FR) context. We achieve superior performance on multiple binding affinity benchmarks through the following contributions:

### 1. CDR-focused supervision yields superior functional representations

CDR-focused models consistently produce the strongest embeddings for downstream binding affinity predictions when evaluated using ridge regression, with *R*^2^ improvements of 20–27% on single-mutant panels and up to 27% on combinatorial libraries using frozen linear probes.

### 2. Paired-only adaptation is sufficient and efficient

Adding large unpaired pretraining or complex curricula that mix unpaired and paired training provides no consistent advantage in our experiments and can introduce drift toward repertoire regularities that are misaligned with VH–VL interface features.

### 3. Compact models can rival larger antibody-specific baselines

The 600M model trained with the same region-aware scheme matches or exceeds larger antibody-specific baselines on multiple benchmarks, indicating that principled supervision can substitute for raw scale.

Our findings parallel those in human language modeling, where smaller domain-adapted models can outperform much larger foundation models on domain-specific tasks [23, 24]. Together, these findings establish a practical framework for antibody PLMs: prioritize paired sequences with CDR-aware supervision over massive pretraining or complex curricula. The best-performing models from this study (AbCDR-ESM2 and AbCDR-ESMC) will be made publicly available for antibody engineering applications.

The remainder of this paper is organized as follows. Section 2 provides comprehensive technical details including datasets, model architectures, masking strategies, training protocols, and compute settings. Section 3 reports region-stratified masked-recovery performance, quantifies the framework–CDR trade-off, and evaluates embedding quality on binding-affinity and combinatorial-mutant benchmarks. Section 4 interprets these findings in the context of antibody representation learning and discusses limitations and future directions.

## 2 Methods

This section describes the datasets, model architectures, training procedures, and evaluation protocols used in this study. Fig. 1 summarizes the overall pipeline, from VH/VL sequence inputs and region-specific masking policies to embedding extraction and linear regression used throughout this study.

**Figure. 1.**
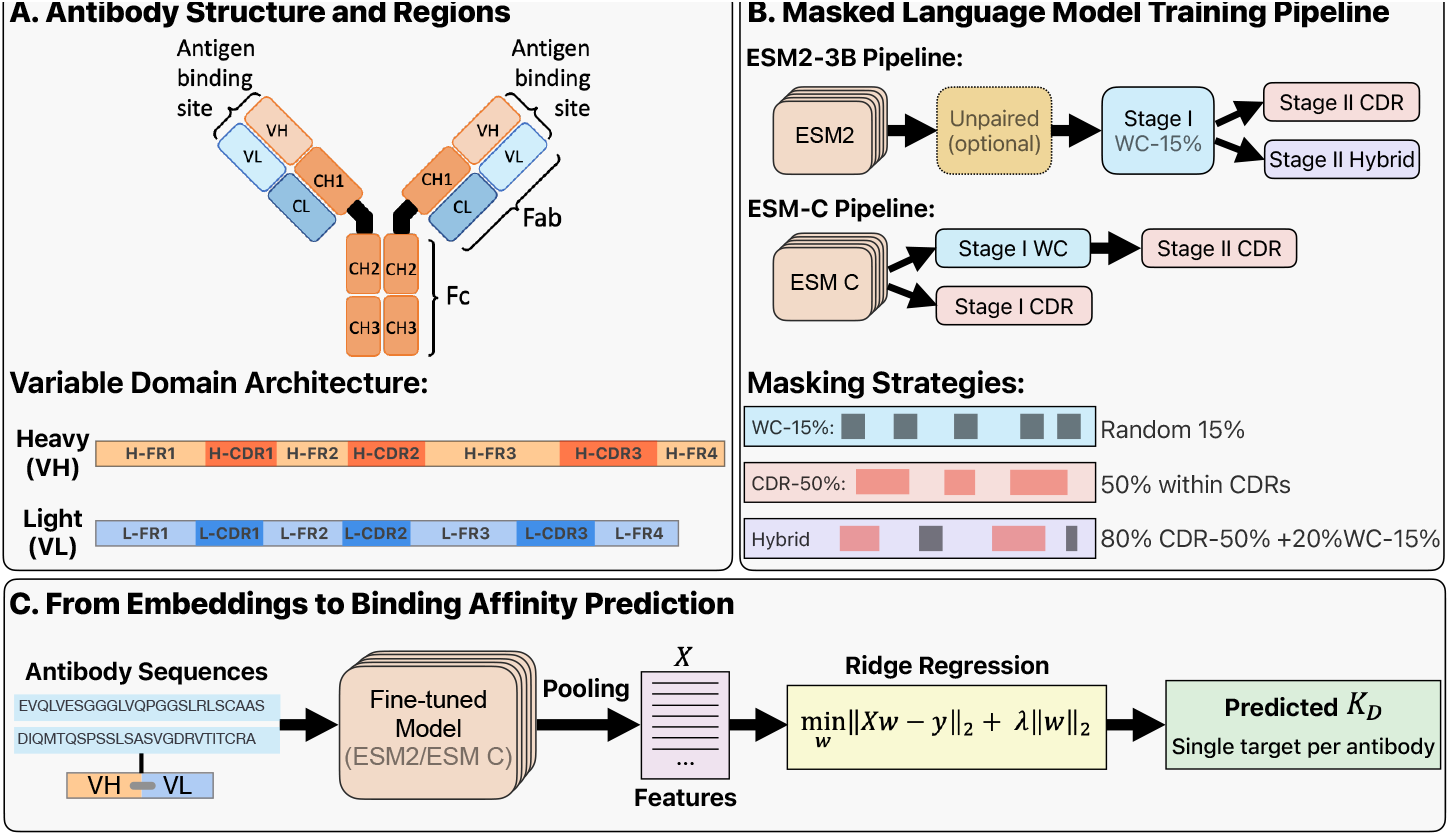
Overview of antibody-specific masked language model training and downstream binding prediction pipeline. **(A)** Antibody structure and region annotation. The Y-shaped antibody comprises paired heavy (VH) and light (VL) variable domains that form the antigen-binding site. Each domain contains four framework regions (FR1–4) providing structural stability and three complementarity-determining regions (CDRs 1–3) that directly contact antigens. **(B)** Two-stage curriculum learning pipelines. ESM2-3B follows an optional unpaired pretraining step, then Stage I whole-chain adaptation, and Stage II refinement with either CDR-focused or hybrid strategies. The 80% and 20% ratios in the hybrid masking indicate the proportion of training samples receiving CDR or WC masking within each batch, respectively. ESM C follows a simpler pipeline with either direct CDR training or a two-stage WC→CDR curriculum. **(C)** Embedding extraction and binding affinity prediction. Fine-tuned models encode paired VH–VL sequences into fixed-dimensional embeddings through mean pooling across all positions. Ridge regression with L2 regularization maps these representations to predicted dissociation constants (*K*_*D*_) for antibody-antigen pairs.

### 2.1 Datasets

#### 2.1.1 Masked Language Modeling

Antibody sequences used for MLM training were collected from the OAS [6, 25], a sequence database of immune repertoires spanning over two billion unpaired sequences and two million paired sequences. Given the focus of the model on antibody property prediction for therapeutics, we removed non-human sequences. Furthermore, we removed potentially autoreactive sequences by excluding repertoires of patients with autoimmune diseases (see Table S2 in supplementary information). To remove near duplicates between training and evaluation, we clustered the remaining antibody sequences at a 95% sequence identity threshold using Linclust for the unpaired pretraining corpus and MMSeqs2 for the paired fine-tuning corpus, both with 80% target coverage (coverage mode 1) and E-value threshold of 0.001. The unpaired pretraining set consists of 1, 220, 072, 064 cluster representatives, from which 12, 200, 720 sequences are selected for validation. For the paired set, 1,617,948 cluster representatives remained, with 20, 225 sequences selected for the validation dataset and another 20, 225 sequences for the test dataset. To enable CDR training, OAS IMGT-labeled CDR sequences were used to delineate sequence indices for masking [26]. The final distributions of residue lengths of the paired set in the variable and CDR regions are shown in Fig. S1.

#### 2.1.2 Downstream datasets

Single mutant antibody datasets were taken from [27–30]. These datasets measured the binding affinity of 2,048, 4,275, and 422 single residue substitution mutations against the target antigens, VEGF, hen egg-white lysozyme, and HER2, respectively. Datasets containing both single and combinatorial mutants were obtained from [31– 33]. These sets provide *K*_*D*_ estimates from high-throughput yeast display for a peptide from the HR2 region of the SARS-CoV-2 spike protein (referred to herein as the anti-HR2 SARS-CoV-2 set), Fluorescein, and H1 Hemagglutinin respectively. Antibody combinatorial mutants were independently verified by calculating average hamming distance between mutants and ensuring it was above the single mutant bound of 2. Unlike the single mutant databases, for the anti-Fluorescein set (found by TiteSeq) and the anti-HR2 SARS-CoV-2 set (found by AlphaSeq), combinatorial mutants were only in the CDR regions while for the anti-H1 Hemagglutinin set (found by MagmaSeq) they were randomly sampled all possible variants between the parent sequence and its universal common ancestor across its 14 changed residue positions. All datasets were downloaded from FLAb [8, 34], except for the anti-HR2 SARS-CoV-2 set which was collected from [35]. Across all datasets, single and combinatorial, affinity data was transformed to log(*K*_*D*_).

### 2.2 Model & training

#### 2.2.1 Model architectures & tokenization

We conducted MLM training using two advanced transformer-based protein language models: ESM2-3B and ESM C (600M). ESM2 is an advanced encoder-only architecture trained on UniRef50 [36] using a masked language modeling objective, and has demonstrated strong generalization across diverse tasks involving protein structure and function prediction. ESM C is a compact encoder trained with increased data and compute resources, intended to improve biological representation learning over ESM2. Although ESM C has only 20% of the parameters of ESM2-3B, it is claimed to offer competitive performance, and we included it in our experiments as a lightweight alternative.

Both models were initialized from their publicly released pretrained checkpoints (hereafter referred to as BASE ESM2 or BASE ESM C) and further adapted using our curated antibody dataset. For tokenization, we used each model’s native tokenizer as released by the authors. Protein sequences were tokenized using a vocabulary of 20 amino acids and special tokens for padding and masking. For paired heavy–light chain modeling, sequences were concatenated with a dedicated separator token (“-”) to preserve their individual identities while enabling inter-chain attention.

#### 2.2.2 Masking policies

We explored three masking regimes. The first follows the conventional whole-chain masking (WC), where 15% of the residues are selected uniformly at random. Among these, 80% are replaced with a special `[MASK]` token, 10% are substituted with a random amino acid token, and 10% are left unchanged, as per the original BERT objective. The second is a domain-informed masking strategy targeting complementarity-determining regions (CDRs), denoted as CDR masking. Here, masking is applied exclusively within the CDR loops, where 50% of CDR residues are masked using the same 80*/*10*/*10 pattern. The 50% masking rate was empirically chosen to match the number of masked residues in the 15% WC scheme, allowing fair comparison. Additionally, we designed a hybrid masking strategy, in which 80% of training samples use CDR masking and 20% use WC masking within each batch, to mitigate the risk of catastrophic forgetting [37, 38] of non-CDR regions. Fig. 1B summarizes the training pipelines for ESM2 and ESM C, which are described in detail below.

#### 2.2.3 Unpaired pretraining (ESM2 only)

To probe whether large unpaired corpora improve subsequent paired adaptation, we pretrain ESM2-3B on unpaired antibody sequences (H or L) using WC-15% for a single epoch. This phase uses a global batch of 512 × 4 × 8 across 32 A100 GPUs (*>* 300 GPU-hours), AdamW with learning rate 1 × 10^−5^, warmup ratio 0.05, and weight decay 0.01. Checkpoints are evaluated every 0.2 epoch; the 0.3-epoch checkpoint initializes downstream paired fine-tuning (ablation). We observed validation loss on the unpaired set continued to improve while paired validation performance peaked early, so we chose the early (0.3) checkpoint to mitigate overfitting to the unpaired distribution.

#### 2.2.4 Paired fine-tuning (ESM2)

We adapt ESM2-3B to paired heavy–light sequences in two stages: *Stage I (WC)* uses whole-chain masking for up to 10 epochs; we evaluate all Stage I epochs and designate epoch 5 as the anchor for Stage II. *Stage II* branches from Stage I WC and is run under two objectives: (i) *CDR* masking at 50% within annotated CDRs; and (ii) *Hybrid* masking with an 80%,CDR / 20%,WC mixture per batch. Stage II runs for up to 10 epochs. To align with downstream performance while avoiding FR degradation from catastrophic forgetting, Stage II checkpoints are selected on validation by a Pareto-knee rule: choose the earliest epoch e^*^ that maximizes CDR(avg) subject to

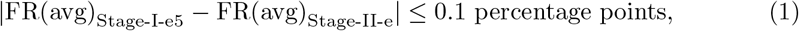

where FR(avg) and CDR(avg) denote mean token-level MLM accuracy within FR and CDR regions, respectively, computed on the paired validation set. This yields e^*^ = 3 for CDR; we also report Hybrid at epoch 3 for consistency. Validation Pareto frontiers are presented in Fig. S2.

#### 2.2.5 Paired fine-tuning (ESM C)

ESM C (600M) is adapted to paired H–L inputs under three regimes: *Stage I (WC-15%)* from BASE; (ii) *Stage I (CDR-50%)* directly from BASE; and (iii) a two-stage *WC-CDR* curriculum that branches from Stage I WC. For parity with ESM2 reporting, we use WC epoch 5 as the Stage II anchor. We apply the same Pareto-knee selection criterion: choose the earliest epoch e^*^ that maximizes CDR(avg) while maintaining framework stability. For Stage I CDR, this yields e^*^ = 4 where the model achieves optimal FR–CDR balance (see Fig. S3). The two-stage curriculum (Stage II) also uses epoch 4 for consistency. We did not adopt a hybrid variant for ESM C, as WC pretraining provided no measurable benefit in our downstream evaluations.

#### 2.2.6 Optimization and compute

All paired fine-tuning runs (ESM2 and ESM C) use identical training settings for fair comparison: mixed-precision (bfloat16) on 16 A100 GPUs, per-device batch size 16 (global batch 256), AdamW (learning rate 2 × 10^−5^), warmup ratio 0.05, weight decay 0.01. Training is implemented with Hugging Face Transformers + Accelerate and DeepSpeed ZeRO.

### 2.3 Masked recovery evaluation

We evaluate paired heavy and light sequences under two masking schemes: WC-15% (uniform over the full chain) and CDR-50% (within annotated CDRs only). Our primary metric is top-1 recovery, defined as the proportion of masked amino acids for which the model’s most probable prediction matches the true residue. We report macro accuracies for frameworks and CDRs, computed as the mean across framework regions (H and L: FR1–FR4) and across CDR regions (H and L: CDR1–CDR3). We also report per-region accuracies for H FR1–FR4, H CDR1–CDR3, L FR1–FR4, and L CDR1–CDR3 (Fig. 1A).

### 2.4 Embedding extraction & downstream heads

To evaluate the learned representations from our antibody-adapted models, we followed the downstream evaluation protocol established in [18] (Fig. 1C). For each paired H–L antibody sequence, we extracted residue-level representations from the final encoder layer. We computed a single embedding vector by averaging across all positions in both heavy and light chains. The resulting embeddings have dimensionality *d* = 2,560 for ESM2, and *d* = 1,152 for ESM C. For baseline model comparisons, paired models (IgBERT and IgT5 [18], AbLang2 [20]) used the same paired averaging as our models, while unpaired models (AntiBERTy [15], AbLang [16], ProtBERT and ProtT5 [39]) concatenated separately-encoded heavy and light chain embeddings following [18].

We employed ridge regression to predict logarithmic dissociation constants (log(*K*_*D*_)) from the extracted embeddings. Binding affinities were log-transformed as *K*_*D*_ values typically span multiple orders of magnitude (*nM* to *mM* range), making linear regression more appropriate in log-space. The regularization strength *λ* was optimized via nested cross-validation: an inner 5-fold cross-validation searched *λ* ∈{1, 10^−1^, …, 10^−6^, 0} to minimize mean squared error on the training folds. The selected *λ* values were 0.1 for ESM2 and 0.001 for ESM C, reflecting the different embedding dimensionalities and model capacities. Each dataset was then reshuffled using a different random seed and 10-fold cross-validation was used for model evaluation.

Coefficient of determination (*R*^2^) was calculated as 1 − *SS*_*res*_*/SS*_*tot*_, measuring the proportion of variance in log *K*_*D*_ explained by the model. Values closer to 1 indicate better predictive performance. Given the inherent noise in binding affinity measurements and the complexity of antibody–antigen interactions, *R*^2^ values in the 0.3–0.4 range represent strong performance for this task.

### 2.5 Statistics and reproducibility

All MLM training runs under the configurations detailed in Section 2.2 utilized fixed random seeds. Masked recovery performance was evaluated on the held-out test set (*n* = 20,225) which was strictly separate from the validation set used for checkpoint selection. To account for stochasticity in the masking process, both validation and test evaluations were performed using two distinct random seeds to generate independent masking patterns. We report region-stratified accuracies averaged across these seeds. Confidence intervals for region-stratified metrics were computed via bootstrap resampling: for each of the two evaluation seeds, we generated 2,000 bootstrap replicates by sampling sequences with replacement, computed per-region accuracies for each replicate, then averaged the corresponding replicates across the two seeds. The 95% confidence intervals represent the 2.5th and 97.5th percentiles of these 2,000 averaged bootstrap replicates.

For downstream embedding evaluations, we employed rigorous repeated crossvalidation to ensure robust statistical estimates. Hyperparameter tuning (regularization strength *λ*) was performed independently within each training fold via the nested 5-fold cross-validation described in Section 2.4 to prevent data leakage. For single-mutant datasets (D44, G6, Trastuzumab), we utilized 10-fold cross-validation, replicating the data splits from [18] for direct benchmarking. For combinatorial mutant datasets (anti-HR2 SARS-CoV-2, anti-Fluorescein, anti-H1 Hemagglutinin), we performed 40 independent random 90%/10% train-test splits. Additionally, for the retrospective analysis of training regimes (Fig. 3), we standardized on 10-fold cross-validation repeated over 40 random seeds per epoch to generate robust mean performance estimates. All reported downstream metrics (*R*^2^, MAE, Spearman’s *ρ*) represent the mean and standard deviation across these cross-validation folds.

## 3 Results

We evaluated our CDR-aware masking strategies (WC, CDR, or hybrid) in two complementary ways. First, we assessed region-stratified masked recovery performance (Section 3.1) to show that CDR-focused training successfully improves representation of these hypervariable regions while maintaining framework accuracy. While these masked recovery metrics demonstrate that our approach learns improved CDR representations, they also serve as proxy measures of overall model quality. As such, we then assessed whether the CDR-focused representation improvements translate to functional utility in the downstream binding affinity prediction task (Section 3.2). This evaluation confirmed that CDR-optimized embeddings substantially improve prediction of antibody–antigen binding across diverse targets, establishing both the validity of our training approach and its practical value for antibody engineering.

### 3.1 Region-stratified masked recovery and trade-offs

#### 3.1.1 ESM2

The base ESM2-3B model shows pronounced performance disparities between framework and CDR regions (Fig. 2A, left), with framework regions achieving 72–92% recovery while CDRs, particularly the hypervariable HCDR3 (35.69%) and LCDR3 (46.06%), perform substantially worse. This gradient reflects the inverse relationship between sequence conservation and functional diversity in antibody recognition.

**Figure. 2.**
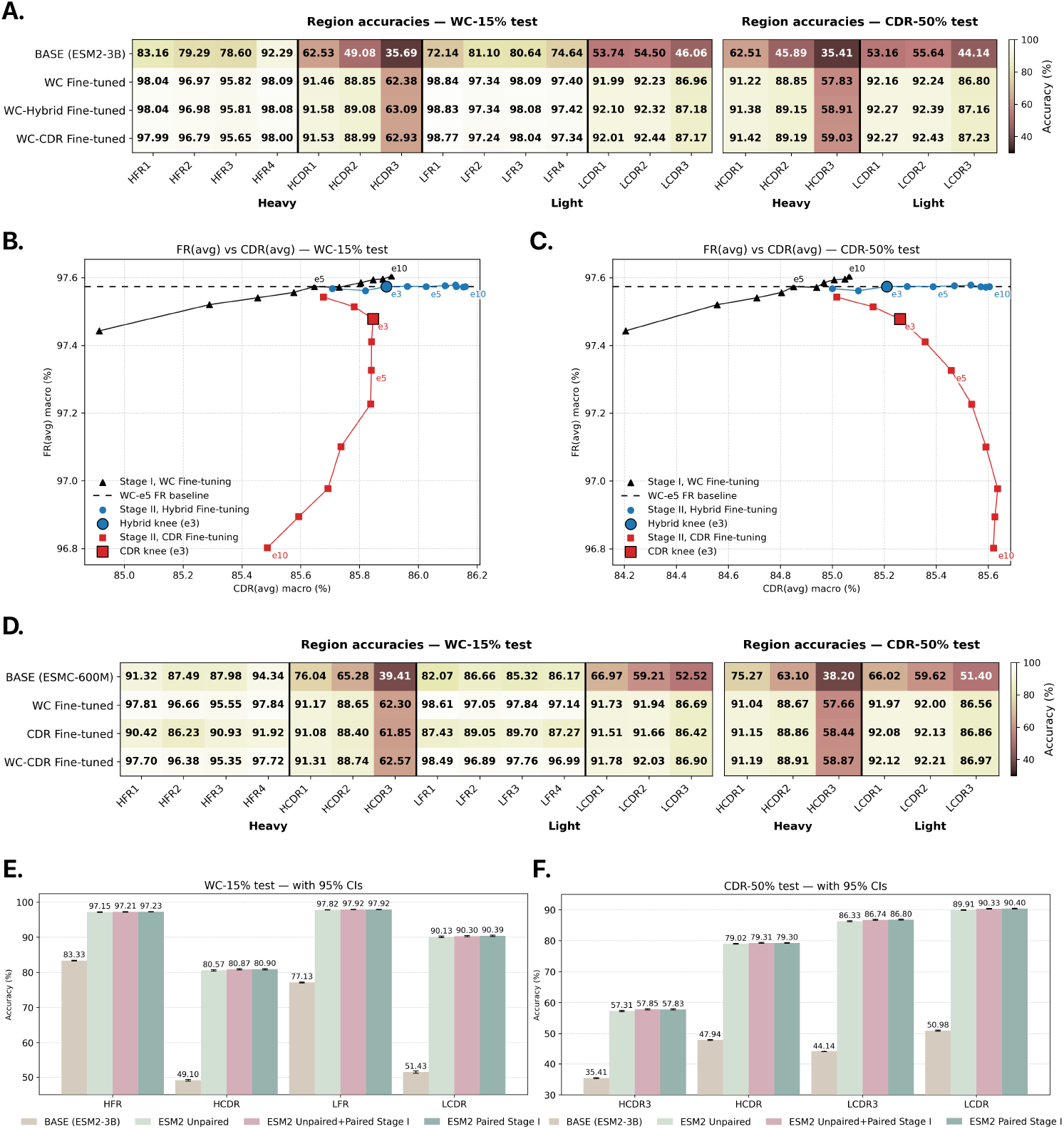
Region-stratified masked recovery. **(A)** ESM2-3B performance across regions. Left: whole-chain masking (WC-15%) test showing per-region top-1 recovery accuracies for base ESM2-3B, Stage I whole-chain fine-tuning (WC, epoch 5), Stage II Hybrid masking (epoch 3), and Stage II CDR-only masking (epoch 3). Right: CDR-specific masking (CDR-50%) test with frameworks unmasked. **(B–C)** Pareto across masking distributions. **(B)** FR(avg) vs. CDR(avg) accuracy on WC-15% test across Stage II training epochs (e = 1–10) starting from Stage I WC (epoch 5, marked by dashed crosshairs). **(C)** Joint evaluation: x-axis shows CDR accuracy on CDR-50% test (frameworks unmasked), y-axis shows FR accuracy on WC-15% test. **(D)** ESM C (600M) performance comparison. Same layout as panel A for ESM C fine-tuned models. **(E–F)** Unpaired pretraining ablation. **(E)** Heavy and light chain framework and CDR macro accuracies on WC-15% test. **(F)** HCDR3, LCDR3, and averaged CDR accuracies on CDR-50% test. Both panels compare Base ESM2, Unpaired-only pretraining, and Unpaired+Paired-WC sequential training, and Stage I WC (paired-only). Bars represent mean accuracy; error bars represent 95% bootstrap confidence intervals. Data represent mean of *n* = 2 independent evaluation seeds (different random masking patterns), with confidence intervals computed from 2,000 bootstrap replicates per seed, resampling from *n* = 20,225 biologically independent test sequences.

**Figure. 3.**
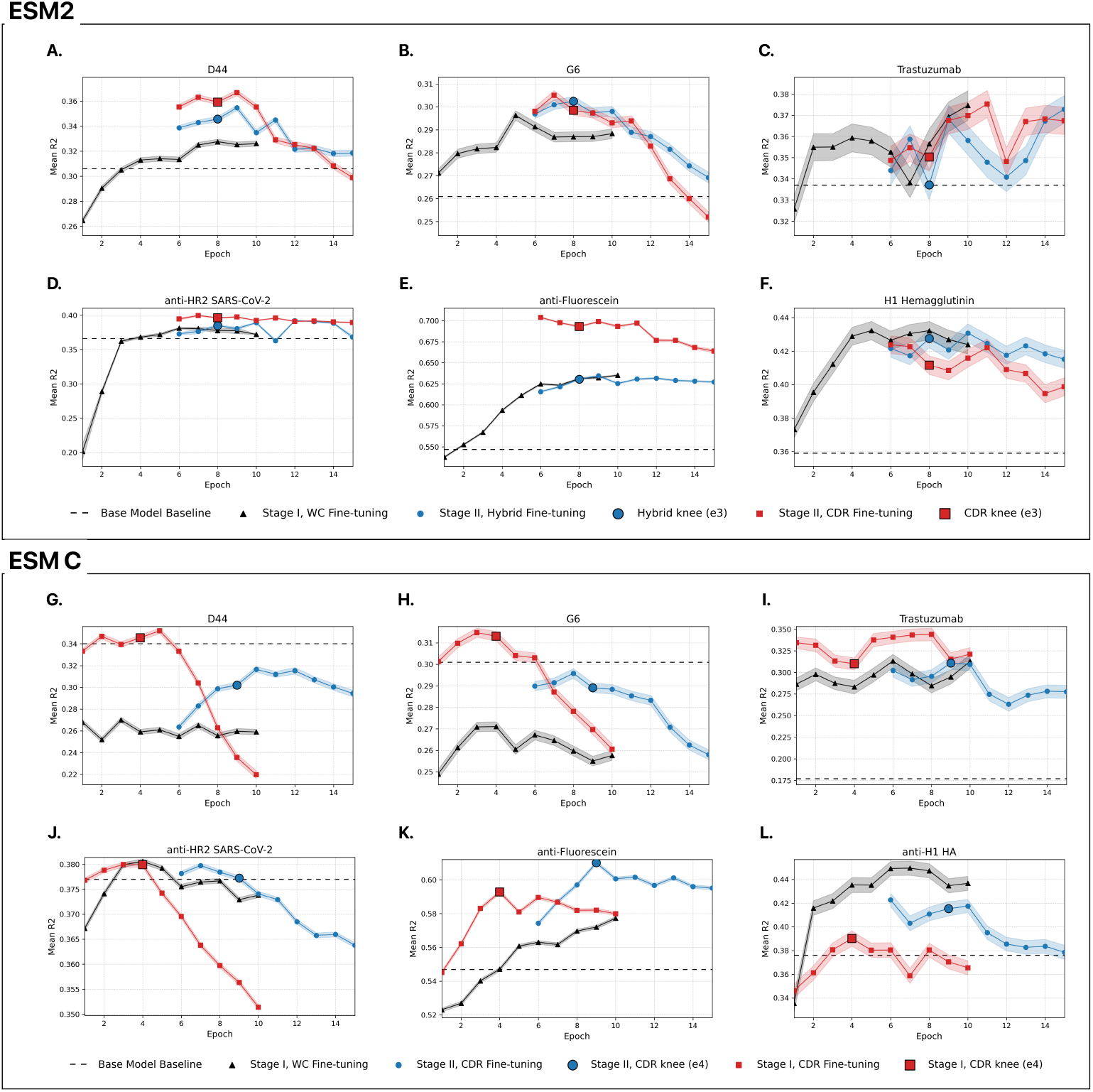
Retrospective analysis of training regimes. *R*^2^ performance of embeddings of each MLM strategy calculated per epoch via 10-fold cross validation split repeated over 40 random seeds per epoch. **(A–F)** ESM2 models on datasets corresponding to single mutant sets (D44, G6, and Trastuzemab) and combinatorial mutant sets (Covid, Fluorescein, H1 Hemagglutinin). **(G–L)** ESM C models on datasets corresponding to single mutant sets (D44, G6, and Trastuzemab) and combinatorial mutant sets (Covid, Fluorescein, H1 Hemagglutinin). Error bands represent the standard error of the mean (SEM) across 40 random seeds.

Whole-chain fine-tuning dramatically improves both regions, with frameworks approaching saturation (97.57% average) and CDRs improving by 35 percentage points (pp) to 85.65%. HCDR3 remains most challenging (62.38%) despite a 75% relative gain. This result is consistent with V(D)J recombination, in which stochastic joining of V, D, and J gene segments generates substantially greater sequence diversity than the V–J joining that gives rise to the other CDRs.

Stage II refinement with CDR-focused or hybrid masking provides additional improvements. Hybrid masking preserves framework performance while improving CDRs (HCDR3: 62.38% →63.09%, LCDR3: 86.96% →87.18%), whereas pure CDR masking achieves similar CDR gains with minimal framework degradation (−0.10 pp). Under CDR-specific evaluation where frameworks provide context (Fig. 2A, right), both Stage II strategies outperform Stage I baseline, with CDR-only masking achieving the highest HCDR3 recovery (59.03% vs. 57.83% for Stage I).

To characterize the framework–CDR trade-off, we analyze Pareto frontiers across Stage II epochs (Fig. 2B–C). Under whole-chain evaluation, the hybrid branch maintains framework stability while improving CDR recovery, whereas CDR-only masking eventually degrades framework representations, reducing the model’s ability to leverage framework context for CDR predictions. Under CDR-specific evaluation (Fig. 2C), Stage II CDR-focused training advances further along the CDR axis when frameworks remain unmasked during evaluation, demonstrating successful specialization.

#### 3.1.2 ESM C

ESM C (600M), despite having only 20% of ESM2’s parameters, demonstrates superior base performance across all regions (Fig. 2D). Framework regions achieve 82–94% recovery and CDRs show marked improvement over ESM2, though HCDR3 remains challenging at 39.41%. WC fine-tuning improves both frameworks (reaching 96–99%) and CDRs (85.41% average), with HCDR3 jumping from 39.41% to 62.30%(+22.89 pp) and LCDR3 going from 52.52% to 86.69% (+34.17 pp). Direct CDR fine-tuning (Stage I CDR) achieves similar CDR performance (85.16% average) but degrades framework accuracy. A two-stage WC–CDR refinement yields only marginal CDR changes while further eroding frameworks, offering no advantage on the WC test. Under CDR-specific evaluation (Fig. 2D, right), CDR-focused fine-tuning shows its intended effect. Stage I CDR surpasses the WC model and performs comparably to Stage II WC→CDR, despite requiring only half the training epochs, suggesting that direct CDR optimization is more efficient than sequential refinement for this architecture.

#### 3.1.3 Unpaired pretraining ablation

We investigated whether large-scale unpaired antibody data could provide a beneficial initialization for paired fine-tuning. Previous work ([18]) has shown that domain-specific pretraining can improve downstream task performance, and the OAS database offers billions of unpaired sequences compared to millions in our paired dataset. We therefore ablated this unpaired pretraining stage by comparing models trained with and without this intermediate step.

We pretrained ESM2-3B on single-chain OAS sequences with WC-15% masking, then fine-tuned on paired sequences. Fig. 2E and 2F compare masked recovery performance with and without this unpaired stage. Unpaired-only pretraining yields a substantial jump over the released base model across all regions, confirming that single-chain antibody data are informative.

However, once we fine-tune on paired data, starting either from the unpaired checkpoint or directly from BASE, the models converge: paired-only WC (Stage I) matches or slightly exceeds the unpaired→paired variant. The unpaired pretraining stage provides no measurable benefit for masked recovery and did not improve downstream binding affinity prediction (Table S1), while adding substantial compute and pipeline complexity.

#### 3.1.4 Comprehensive comparison

We compare our Stage II models against published antibody-specific baselines (IgBERT, IgT5) on masked recovery. While direct cross-paper comparisons are approximate due to different data splits, both studies use similarly sized test sets (~ 20k sequences) from the same OAS source, enabling indicative comparisons.

On the WC-15% test (Table 1), our models achieve performance comparable to IgBERT and IgT5 across most regions, with consistent improvements in HCDR3 and LCDR3 in all our models. Given potential data split differences, we interpret these results as demonstrating that CDR-focused fine-tuning from general protein models can match the masked recovery performance of antibody-specific architectures trained on massive unpaired corpora, while requiring substantially less computational resources (8 epochs vs. 46 epochs for IgBERT).

**Table 1.** Region accuracies on WC and CDR masked test. Top-1 recovery (%) on a held-out test set of *n* = 20,225 paired heavy–light sequences. “Total VH/VL” are token-weighted micro averages over all masked heavy/light tokens. Published baselines (IgBert, IgT5) report WC-15% results on ~20k test sets from their original splits; splits may differ from ours, so cross-paper comparisons are indicative rather than strictly matched. Total HCDR/LCDR are micro-averages over masked CDR tokens in heavy/light chains. Bold numbers indicate the highest accuracy within each region.

| Heavy-chain regions (WC-15% test) |  |  |  |  |  |  |  |  |
| --- | --- | --- | --- | --- | --- | --- | --- | --- |
| Model | HFR1 | HFR2 | HFR3 | HFR4 | HCDR1 | HCDR2 | HCDR3 | Total VH |
| *Base Models* |  |  |  |  |  |  |  |  |
| Base ESM2-3B | 83.16 | 79.29 | 78.60 | 92.29 | 62.53 | 49.08 | 35.69 | 72.32 |
| Base ESMC-600M | 91.32 | 87.49 | 87.98 | 94.34 | 76.04 | 65.28 | 39.41 | 80.63 |
| *Published Model Results* |  |  |  |  |  |  |  |  |
| IgBert Unpaired | 97.91 | 96.55 | 95.52 | 97.98 | 90.43 | 88.41 | 59.24 | 90.99 |
| IgT5 Unpaired | 97.90 | 96.71 | 95.60 | 98.25 | 90.92 | 88.39 | 60.35 | 91.21 |
| IgBert | 98.10 | 96.90 | 95.76 | 98.09 | 91.30 | 88.65 | 60.12 | 91.29 |
| IgT5 | **98.20** | 96.87 | 95.74 | **98.28** | 91.50 | **89.36** | 61.96 | 91.63 |
| *Our Models - ESM2* |  |  |  |  |  |  |  |  |
| ESM2 Unpaired | 97.99 | 96.87 | 95.72 | 98.03 | 91.21 | 88.71 | 61.78 | 91.42 |
| ESM2 Stage I WC (e5) | 98.04 | 96.97 | **95.82** | 98.09 | 91.46 | 88.85 | 62.38 | 91.58 |
| ESM2 Stage II Hybrid (e3) | 98.04 | **96.98** | 95.81 | 98.08 | **91.58** | 89.08 | **63.09** | **91.69** |
| ESM2 Stage II CDR (e3) | 97.99 | 96.79 | 95.65 | 98.00 | 91.53 | 88.99 | 62.93 | 91.57 |
| Light-chain regions (WC-15% test) |  |  |  |  |  |  |  |  |
| Model | LFR1 | LFR2 | LFR3 | LFR4 | LCDR1 | LCDR2 | LCDR3 | Total VL |
| *Base Models* |  |  |  |  |  |  |  |  |
| Base ESM2-3B | 72.14 | 81.10 | 80.64 | 74.64 | 53.74 | 54.50 | 46.06 | 72.45 |
| Base ESMC-600M | 82.07 | 86.66 | 85.32 | 86.17 | 66.97 | 59.21 | 52.52 | 79.88 |
| *Published Model Results* |  |  |  |  |  |  |  |  |
| IgBert Unpaired | 98.04 | 97.04 | 97.39 | 96.56 | 90.81 | 89.85 | 84.61 | 95.60 |
| IgT5 Unpaired | 98.09 | 96.75 | 97.52 | 91.71 | 90.76 | 90.93 | 84.23 | 95.15 |
| IgBert | **98.85** | **97.38** | 98.07 | 97.40 | **92.32** | 91.49 | 86.34 | 96.47 |
| IgT5 | 98.78 | 97.35 | **98.15** | **97.84** | 92.22 | 91.63 | 86.93 | **96.56** |
| *Our Models - ESM2* |  |  |  |  |  |  |  |  |
| ESM2 Unpaired | 98.73 | 97.30 | 98.05 | 97.21 | 92.14 | 91.58 | 86.67 | 96.39 |
| ESM2 Stage I WC (e5) | 98.84 | 97.34 | 98.09 | 97.40 | 91.99 | 92.23 | 86.96 | 96.51 |
| ESM2 Stage II Hybrid (e3) | 98.83 | 97.34 | 98.08 | 97.42 | 92.10 | 92.32 | **87.18** | 96.54 |
| ESM2 Stage II CDR (e3) | 98.77 | 97.24 | 98.04 | 97.34 | 92.01 | **92.44** | 87.17 | 96.48 |
| CDR regions (CDR-50% test) |  |  |  |  |  |  |  |  |
| Model | HCDR1 | HCDR2 | HCDR3 | Total HCDR | LCDR1 | LCDR2 | LCDR3 | Total LCDR |
| *Base Models* |  |  |  |  |  |  |  |  |
| Base ESM2-3B | 62.51 | 45.89 | 35.41 | 45.06 | 53.16 | 55.64 | 44.14 | 49.21 |
| Base ESMC-600M | 75.27 | 63.10 | 38.20 | 53.98 | 66.02 | 59.62 | 51.40 | 58.04 |
| *Results from Published Models<sup>1</sup>* |  |  |  |  |  |  |  |  |
| IgBERT Unpaired | 91.03 | 88.68 | 57.53 | 73.90 | 91.98 | 91.40 | 86.25 | 89.14 |
| IgBert | 91.35 | 89.07 | 58.44 | 74.53 | **92.27** | 92.34 | 86.92 | 89.71 |
| *Our Models - ESM2* |  |  |  |  |  |  |  |  |
| ESM2 Unpaired | 91.08 | 88.66 | 57.31 | 73.80 | 92.08 | 91.33 | 86.33 | 89.21 |
| ESM2 Stage I WC (e5) | 91.22 | 88.85 | 57.83 | 74.14 | 92.16 | 92.24 | 86.80 | 89.60 |
| ESM2 Stage II Hybrid (e3) | 91.38 | 89.15 | 58.91 | 74.79 | **92.27** | 92.39 | 87.16 | 89.84 |
| ESM2 Stage II CDR (e3) | **91.42** | **89.19** | **59.03** | **74.87** | **92.27** | **92.43** | **87.23** | **89.88** |

<sup>1</sup> Published baselines are not released with CDR-50% evaluation; we re-evaluated these models under the CDR-50% policy on our test split. Because our split may overlap their training data, the reported values for these baselines could be optimistic relative to a fully disjoint test.

For the CDR-50% test (Table 1, bottom), we evaluated published baselines on our test split since this evaluation protocol was not included in their original work. Because our test split may overlap with their training data, the reported baseline values represent an upper bound on their performance under this evaluation. Despite this potential advantage, our Stage II CDR models achieve the highest performance across all CDR regions (74.87% HCDR, 89.88% LCDR), demonstrating effective CDR specialization.

Importantly, these masked recovery results motivated examining downstream functional prediction (Section 3.2), where CDR-optimized representations provide the strongest evidence of improved antibody modeling through substantial gains in binding affinity prediction.

### 3.2 Embedding quality & transfer to antibody binding affinity prediction

To assess whether this region-stratified masking pipeline leads to improved functional prediction, we selected models for downstream embedding evaluation based on validation set performance. We chose checkpoints that: (a) achieved optimal masked CDR recovery, (b) demonstrated Pareto optimality between FR and CDR recovery, and (c) represented the earliest epoch meeting these criteria to prevent overfitting to the masked language modeling objective. The selected models were ESM2 Stage II CDR epoch 3 and ESM C Stage I CDR epoch 4, hereafter referred to as AbCDR-ESM2 and AbCDR-ESMC, respectively.

#### 3.2.1 CDR-focused training improves binding affinity prediction

The embedding quality of AbCDR-ESM2 and AbCDR-ESMC was evaluated on three therapeutic antibody binding datasets (Table 2) from FLAb [8], where each dataset measured binding affinities of single residue mutants. We follow the evaluation protocol established by [18] and compare against their reported baseline performance. Our reproduced AntiBERTy and AbLang results closely matched these published values, confirming the reliability of our evaluation protocol. We additionally benchmarked AbLang2 [20], which was not previously evaluated on these datasets.

**Table 2.**
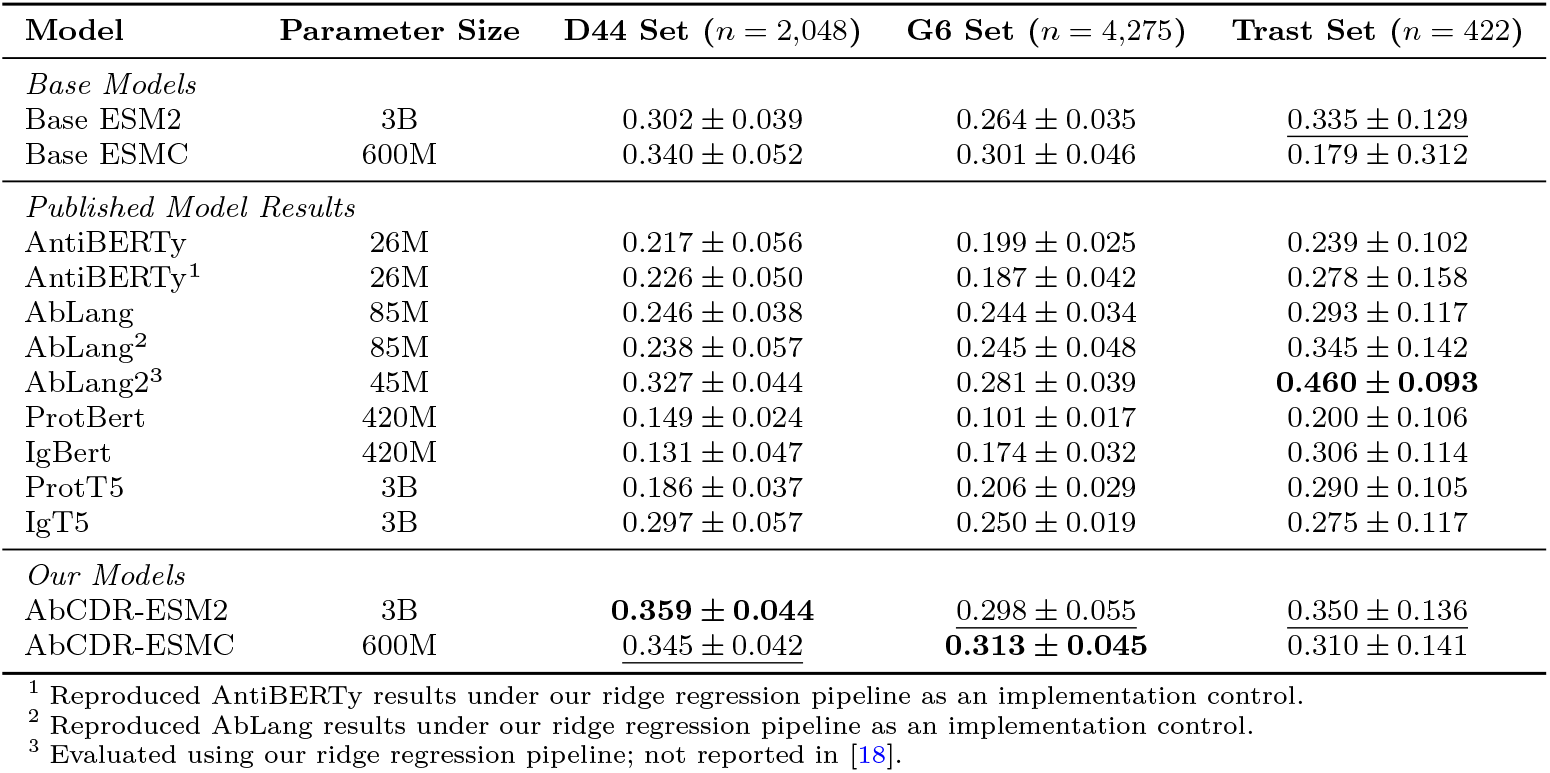
Comparative performance of embeddings binding regression *R*^2^. Values represent mean *R*^2^ ± standard deviation calculated using 10-fold cross-validation replicating original splits from [18], enabling cross-paper comparison. Published model results are taken directly from [18] unless indicated otherwise. First and second best performing models for each metric indicated in bold and underline, respectively.

Compared to their base counterparts, our antibody-adapted models consistently outperformed across all benchmarks; ESM2 adaptation improved *R*^2^ from 0.302 to 0.359 on D44 (+18.8%), from 0.264 to 0.298 on G6 (+12.8%), and modestly increased Trast (0.335 → 0.350; +4.4%).

The compact AbCDR-ESMC model was highly efficient: targeted adaptation yielded the second highest *R*^2^ on D44 (0.345) and highest *R*^2^ on G6 (0.313), surpassing the 3B-parameter IgT5 by 16% and 25%, respectively, despite using 5x fewer parameters and only paired fine-tuning. This validates that domain-specific fine-tuning benefits models across different scales and architectures.

Compared to other antibody-specific models, our approach achieved superior performance without massive unpaired pretraining. While IgBert, AbLang2, and IgT5 leverage billions of unpaired sequences, our adapted models attained 20–170% higher *R*^2^ with just 8 epochs of paired training for ESM2 (5 WC + 3 CDR) or 4 epochs of direct CDR training for ESM C. Notably, AbLang2 achieved the highest *R*^2^ on Trast (0.460), the only case in which a published model surpassed our adapted models. However, Trast is also the smallest dataset (*n* = 422) with the highest cross-validation variance.

Interestingly, base ESM2 already matched or exceeded IgT5 across datasets, and base ESM C did so on D44 and G6 (but not Trast). The consistent gains from CDR-focused fine-tuning, ranging from ~ 4–20% for ESM2 and up to 73% for ESM C on Trast, indicate that strategic masking on high-quality paired data provides benefits orthogonal to model scale.

#### 3.2.2 Generalization to combinatorial mutant prediction

To evaluate generalization beyond single mutant binding affinities, we extended our evaluation of AbCDR-ESM2 and AbCDR-ESMC to include datasets with both single and combinatorial antibody mutants across three diverse antigen systems (Table 3). We evaluated all baseline models using the embedding extraction and ridge regression protocol described in Section 2.4.

**Table 3.** Downstream regression prediction of free energies of combinatorial mutants using a linear model. All models were evaluated using the same embedding extraction and ridge regression protocol. Performance metrics represent mean ± standard deviation across 40 independent random 90%*/*10% train-test splits. First and second best performing models for each metric indicated in bold and underline, respectively.

| anti-HR2 SARS-CoV-2 Set ($n = 71,830$) | | | | |
| --- | --- | --- | --- | --- |
| Model | Parameter Size | MAE | Mean $R^2$ | Spearman's $\rho$ |
| *Base Models* |  |  |  |  |
| Base ESM2 | 3B | $0.865 \pm 0.025$ | $0.366 \pm 0.036$ | $0.580 \pm 0.022$ |
| Base ESMC | 600M | $0.857 \pm 0.007$ | $0.377 \pm 0.010$ | $0.587 \pm 0.008$ |
| *Results from Published Models* |  |  |  |  |
| AntiBERTy | 26M | $0.917 \pm 0.007$ | $0.297 \pm 0.010$ | $0.521 \pm 0.009$ |
| AbLang | 85M | $0.895 \pm 0.008$ | $0.328 \pm 0.010$ | $0.551 \pm 0.009$ |
| AbLang2 | 45M | $0.896 \pm 0.008$ | $0.324 \pm 0.010$ | $0.546 \pm 0.009$ |
| ProtBert | 420M | $0.879 \pm 0.007$ | $0.351 \pm 0.010$ | $0.566 \pm 0.009$ |
| IgBert | 420M | $0.882 \pm 0.078$ | $0.338 \pm 0.178$ | $0.563 \pm 0.034$ |
| ProtT5 | 3B | $0.872 \pm 0.008$ | $0.359 \pm 0.009$ | $0.575 \pm 0.008$ |
| IgT5 | 3B | $0.866 \pm 0.007$ | $0.364 \pm 0.009$ | $0.577 \pm 0.008$ |
| *Our Models* |  |  |  |  |
| AbCDR-ESM2 | 3B | **$0.841 \pm 0.008$** | **$0.396 \pm 0.014$** | **$0.599 \pm 0.013$** |
| AbCDR-ESMC | 600M | <u>$0.855 \pm 0.007$</u> | <u>$0.379 \pm 0.009$</u> | <u>$0.587 \pm 0.008$</u> |
| anti-Fluorescein Set ($n = 11,052$) | | | | |
| Model | Parameter Size | MAE | Mean $R^2$ | Spearman's $\rho$ |
| *Base Models* |  |  |  |  |
| Base ESM2 | 3B | $0.791 \pm 0.016$ | $0.547 \pm 0.019$ | $0.697 \pm 0.017$ |
| Base ESMC | 600M | $0.788 \pm 0.034$ | $0.547 \pm 0.057$ | $0.694 \pm 0.022$ |
| *Results from Published Models* |  |  |  |  |
| AntiBERTy | 26M | $0.853 \pm 0.018$ | $0.464 \pm 0.023$ | $0.648 \pm 0.019$ |
| AbLang | 85M | $0.809 \pm 0.018$ | $0.531 \pm 0.023$ | $0.686 \pm 0.018$ |
| AbLang2 | 45M | $0.888 \pm 0.018$ | $0.446 \pm 0.023$ | $0.659 \pm 0.018$ |
| ProtBert | 420M | $0.920 \pm 0.020$ | $0.410 \pm 0.016$ | $0.639 \pm 0.021$ |
| IgBert | 420M | $0.812 \pm 0.017$ | $0.517 \pm 0.023$ | $0.681 \pm 0.017$ |
| ProtT5 | 3B | $0.877 \pm 0.018$ | $0.464 \pm 0.023$ | $0.673 \pm 0.019$ |
| IgT5 | 3B | $0.872 \pm 0.017$ | $0.459 \pm 0.024$ | $0.664 \pm 0.018$ |
| *Our Models* |  |  |  |  |
| AbCDR-ESM2 | 3B | **$0.594 \pm 0.014$** | **$0.693 \pm 0.025$** | **$0.730 \pm 0.044$** |
| AbCDR-ESMC | 600M | <u>$0.737 \pm 0.015$</u> | <u>$0.592 \pm 0.021$</u> | <u>$0.705 \pm 0.017$</u> |
| anti-H1 HA Set ($n = 1,038$) | | | | |
| Model | Parameter Size | MAE | Mean $R^2$ | Spearman's $\rho$ |
| *Base Models* |  |  |  |  |
| Base ESM2 | 3B | $0.168 \pm 0.019$ | $0.359 \pm 0.100$ | $0.721 \pm 0.051$ |
| Base ESMC | 600M | $0.160 \pm 0.019$ | $0.376 \pm 0.124$ | $0.756 \pm 0.047$ |
| *Results from Published Models* |  |  |  |  |
| AntiBERTy | 26M | $0.166 \pm 0.019$ | $0.355 \pm 0.123$ | $0.738 \pm 0.050$ |
| AbLang | 85M | $0.170 \pm 0.021$ | $0.291 \pm 0.145$ | $0.720 \pm 0.056$ |
| AbLang2 | 45M | $0.161 \pm 0.020$ | $0.379 \pm 0.110$ | $0.753 \pm 0.046$ |
| ProtBert | 420M | $0.188 \pm 0.017$ | $0.269 \pm 0.128$ | $0.675 \pm 0.100$ |
| IgBert | 420M | $0.161 \pm 0.021$ | $0.376 \pm 0.129$ | $0.742 \pm 0.052$ |
| ProtT5 | 3B | $0.178 \pm 0.019$ | $0.331 \pm 0.096$ | $0.704 \pm 0.054$ |
| IgT5 | 3B | $0.164 \pm 0.019$ | $0.370 \pm 0.102$ | $0.735 \pm 0.049$ |
| *Our Models* |  |  |  |  |
| AbCDR-ESM2 | 3B | **$0.155 \pm 0.019$** | **$0.411 \pm 0.113$** | **$0.758 \pm 0.048$** |
| AbCDR-ESMC | 600M | <u>$0.156 \pm 0.019$</u> | <u>$0.390 \pm 0.132$</u> | <u>$0.758 \pm 0.050$</u> |

Our adapted models demonstrated consistent improvements across all datasets, with particularly striking gains on the anti-Fluorescein set where AbCDR-ESM2 improved *R*^2^ from 0.547 to 0.693, a 26.6% relative increase and a 30.5% relative increase compared to the best published model, AbLang. This dataset’s focus on multisite CDR mutations may particularly benefit from our CDR-focused training, as these higher-order CDR correlations are learned during 50% CDR masking.

On the large-scale anti-HR2 SARS-CoV-2 dataset (*n* = 71,830), AbCDR-ESM2 achieved *R*^2^ = 0.396, surpassing the best published model, IgT5, by 8.8% while reducing MAE from 0.865 to 0.841. ESM C showed more modest gains, likely because its base performance already exceeded all published baselines. This pattern (stronger base models showing smaller but consistent improvements) recurred across datasets, suggesting that our adaptation provides orthogonal benefits to architectural advances.

For H1 Hemagglutinin, adapted AbCDR-ESM2 achieved the highest *R*^2^ (0.411) but had an almost equivalent Spearman correlation (0.758) to AbCDR-ESMC. The consistent improvements across antigens with different binding modes (HR2 region of SARS-CoV-2 spike protein, small molecule fluorescein, hemagglutinin), both with respect to the base model and leading published models, demonstrate that the two stage CDR-focused training captures generalizable patterns rather than target-specific biases.

#### 3.2.3 Comprehensive embedding comparison

AbCDR-ESM2 and AbCDR-ESMC were selected as the best ESM2 and ESM C based models in an unsupervised fashion: based on their superior masked CDR recovery while being Pareto optimal with respect to FR recovery. We investigated this model selection strategy retrospectively and systematically compared it to downstream embedding performances of other MLM strategies. To generate a robust estimate of the mean performance of each strategy, 10-fold cross validation performance was averaged over 40 random seeds per epoch.

The plots in Fig. 3 show modest agreement across datasets between the selection of Stage II CDR epoch 3 for ESM2 (AbCDR-ESM2) and Stage I CDR epoch 4 for ESM C (AbCDR-ESMC) and the maximum embedding performance across epochs. This validates our Pareto approach as a sensible unsupervised strategy for model selection. In particular, this helps avoid observed embedding degeneration in later epochs of training, as the dilution of broad sequence level features in favor of CDR features begins to worsen downstream embeddings.

We also confirmed that the benefit of CDR masking is independent of training time and mutational distance. AbCDR-ESM2 generated a relative improvement of 0.76%, 9.65%, 9.12%, and 3.98% over the best epoch of WC model for the single mutant sets of G6, D44 and the combinatorial mutant sets of anti-Fluorescein, and anti-HR2 SARS-CoV-2, respectively. Similarly, AbCDR-ESM2 outperforms Stage II Hybrid for three datasets with matching performance, within standard error, for the remaining three.

For AbCDR-ESMC, we observed a more pronounced effect; in some cases, involvement of WC masking (D44 and G6) led to worse performance than the base model. AbCDR-ESMC led to a relative gain 16.1%, 25.7%, and 1.0% over the best epoch of WC masking for G6, D44, and anti-Fluorescein respectively, with matching performance for anti-HR2 SARS-CoV-2 and Trast.

Notably, there are dataset exceptions, as WC masking outperforms any CDR masking strategy for anti-H1 HA while Trast shows no difference in any masking strategy. However, these sets are noticeably smaller (*n* = 1,038 and *n* = 422), which limits confidence whether the observed mean reflects the true underlying distribution for these two antibodies. Aside from anti-H1 HA, CDR masking strategies, from either base ESM models, achieved superior binding prediction compared to traditional WC masking strategies.

## 4 Discussion

Our ablations indicate that paired VH–VL fine-tuning is sufficient and often preferable for learning antibody representations. With standard whole-chain masking, pairedonly adaptation matched or exceeded variants that first underwent large-scale unpaired pretraining on masked-recovery objectives (Fig. 2E and 2F), consistent with the central role of CDR-mediated paratopes and the VH–VL interface in antigen recognition, signals that single-chain objectives cannot fully capture [40–42]. Although unpaired pretraining improved token recovery, it did not translate into downstream binding gains in our trials, suggesting representation drift: single-chain optimization can over-weight framework regularities and germline/frequency biases that are weakly predictive of pair-dependent binding [43, 44]. Practically, small but high-quality paired sets provided sufficient cross-chain signal for affinity transfer, whereas additional unpaired pretraining risked overfitting to repertoire statistics misaligned with functional variation.

A similar pattern appears in published antibody-specific PLMs trained on extensive unpaired corpora: for IgBERT and IgT5, the unpaired → paired pipeline does not yield monotonic improvements across benchmarks. On several datasets the base or unpaired checkpoints are comparable to, or better than, their paired-adapted counterparts (see Table 2; cf. [18]). This lack of a consistent base → unpaired → paired improvement curve supports the view that mixing objectives and data regimes can misalign the embedding space unless supervision explicitly preserves VH–VL interaction features. By contrast, our paired-only adaptation (and, subsequently, CDR-focused supervision) yields reliable gains without massive unpaired antibody pretraining, reinforcing a simple takeaway: when evaluation targets pair-dependent biophysics, training signals that respect the heavy–light joint distribution matter more than scale.

Beyond the efficiency of paired-only training, our findings align with and extend recent work showing that shifting MLM signal toward high-entropy, binding-relevant residues (e.g., via preferential masking of CDR regions or non-germline-weighted losses) improves antibody representations [20, 21]. Here, we show that this effect persists for paired VH–VL models and translates into robust binding-affinity gains, even with modest framework recovery trade-offs. This pattern held across both architectures: ESM2 improved binding *R*^2^ by up to 27% on combinatorial mutant prediction (Fluorescein dataset), while the compact ESM C achieved state-of-the-art performance across multiple benchmarks.

Why would models that perform worse at predicting framework residues excel at binding prediction? The answer illuminates a core principle of antibody function. Frameworks provide structural scaffolding but vary little across antibodies, making them easy to predict yet uninformative for binding specificity. CDRs, by contrast, harbor the sequence diversity that enables antigen recognition. When models train on uniform masking across all positions, they allocate capacity to “solving” predictable framework positions rather than learning the complex patterns within CDRs. Our CDR-focused approach redirects this capacity where it matters most: forcing models to capture subtle covariation within and between CDR loops that determine how antibodies recognize their targets.

The Pareto analysis (Fig. 2B and 2C) captures this trade-off quantitatively. Pure CDR masking drives models toward better CDR representations even as framework accuracy plateaus, while hybrid approaches struggle to balance both objectives. That these CDR-optimized models generalize across diverse antigens, from viral spike proteins to small-molecule fluorescein, confirms they learn fundamental recognition principles rather than dataset-specific patterns. In essence, by aligning training objectives with biological function (CDRs determine specificity), we achieve representations that better predict what ultimately matters: whether an antibody will bind its target.

While our CDR-focused masking strategy demonstrates clear advantages for binding affinity prediction, several limitations merit consideration. First, we evaluated only ESM-family architectures; the generalizability to other model families (e.g., ProtT5, IgBERT) with different positional encodings remains unexplored. Second, our assessment focused exclusively on binding prediction; whether CDR-focused representations improve other therapeutic properties such as stability, expression, or immunogenicity requires investigation. The observed trade-off between CDR and framework recovery may have different implications for these properties.

Despite these limitations, our work establishes that biologically-motivated training strategies can achieve state-of-the-art performance with remarkable computational efficiency and opens a direction for future research to investigate and optimize masking rates within the parameter space. These findings provide a practical framework for developing next-generation antibody language models that prioritize functional relevance over scale.

## 5 Data availability

All source data underlying the figures and tables in this manuscript are provided as Supplementary Data in CSV format.

Raw antibody sequences were obtained from the Observed Antibody Space (OAS) [6, 25], accessible at [25]. Raw binding affinity datasets were obtained from [34, 35] and from published sources [27–33].

The processed and curated datasets generated and used in this study, including clustered antibody sequences with train/validation/test splits and IMGT CDR annotations, are publicly available at Zenodo under the Creative Commons Attribution 4.0 International (CC-BY 4.0) license and can be accessed via DOI: 10.5281/zenodo.18762309 [45]. Curated binding affinity evaluation datasets are similarly publicly available at Zenodo under the CC-BY 4.0 license via DOI: 10.5281/zenodo.18762978 [46].

## 6 Code Availability

The custom code used to develop the models, perform the analyses, and generate results in this study is publicly available and has been deposited in GitHub at https://github.com/noc-lab/AbCDR-ESM, under the MIT license. The specific version of the code associated with this publication is archived in Zenodo and is accessible via DOI: 10.5281/zenodo.19900188 [47].

Additionally, all original pretrained model checkpoints (AbCDR-ESM2 and AbCDR-ESMC) developed in this study are publicly available under the MIT license without access restrictions. They are deposited on the HuggingFace Hub and can be accessed at https://huggingface.co/Paschalidis-NOC-Lab/AbCDR-ESM2 and https://huggingface.co/Paschalidis-NOC-Lab/AbCDR-ESMC. Full instructions for environment setup, code execution, and model inference are provided in the repository documentation.

## Author Contributions

M.T. and K.C.W. carried out the design of the computational experiments, implementation, analysis of the results, and writing of the manuscript. B.H. contributed code, E.J. curated the data sets, and Y.J. performed calculations. D.K., J.M., and S.V. contributed to the analysis of the results and editing of the paper. I.Ch. P. and DJ-M conceptualized the work, oversaw the project, and finalized the manuscript. DJ-M obtained the funding.

