## Supplementary Tables and FIgures for "Preferential CDR masking in paired antibody language models improves binding affinity prediction"

### 1 Supplementary Tables and Figures

**Table S1 Downstream embedding performance after unpaired MLM training.** Anti-HR2 SARS-CoV-2 calculated at random 10/90 split due to computational demand; otherwise mean performance calculated via 10-fold cross validation with a single seed. Best performing models for a given metric indicated in bold.

| Model | D44 | | GP6 | | Trastuzumab | | anti-HR2 SARS-CoV-2 | |
|---|---|---|---|---|---|---|---|---|
| | $R^2$ | Spearman's $\rho$ | $R^2$ | Spearman's $\rho$ | $R^2$ | Spearman's $\rho$ | $R^2$ | Spearman's $\rho$ |
| Base ESM2 | **0.298 ± 0.054** | **0.541 ± 0.073** | **0.261 ± 0.047** | **0.505 ± 0.029** | 0.337 ± 0.079 | 0.595 ± 0.087 | 0.351 | **0.563** |
| ESM2 Unpaired Ab Model | 0.284 ± 0.050 | 0.528 ± 0.066 | 0.253 ± 0.049 | 0.500 ± 0.035 | **0.397 ± 0.109** | **0.651 ± 0.071** | **0.352** | 0.562 |

**Table S2 Removed patient sequence repertoires.** Repertoires from patients with autoimmune diseases were removed to focus on therapeutic antibody prediction.

| Dataset | Removed Disease |
|---|---|
| Unpaired | Multiple Sclerosis<br>Light Chain Amyloidosis<br>Chronic Lymphocytic Leukemia<br>Celiac<br>POEMS Syndrome<br>Systemic Lupus Erythematosus |
| Paired | Multiple Sclerosis |

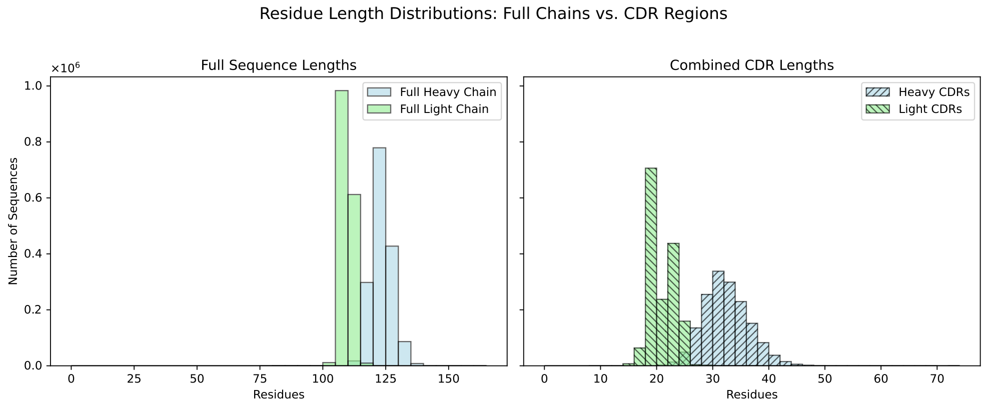


**Fig. S1 Residue Length Distributions.** Final distributions of residue lengths of the paired set in the variable and CDR regions.

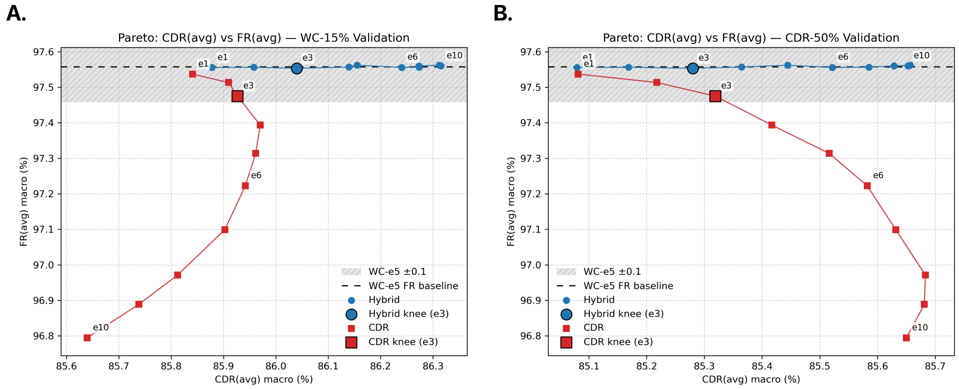


**Fig. S2 ESM2 Validation Pareto. (A)** FR(avg) vs. CDR(avg) accuracy on WC-15% validation across Stage II training epochs (e = 1–10) starting from Stage I WC (epoch 5, marked by dashed crosshairs). **(B)** Joint evaluation: x-axis shows CDR accuracy on CDR-50% validation (frameworks unmasked), y-axis shows FR accuracy on WC-15% validation. The earliest epoch maximizing CDR(avg) subject to 0.1 framework drift is epoch 3 and is therefore chosen for the subsequent analysis.

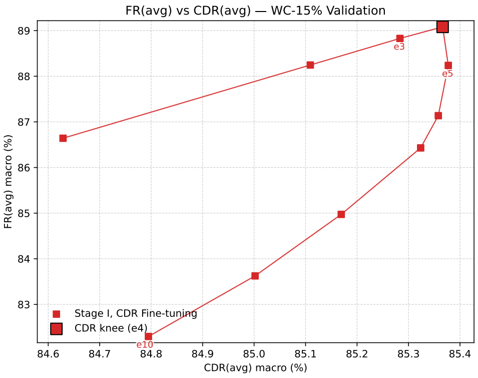


**Fig. S3 ESM C Validation Pareto.** Pareto frontier for ESM C model showing framework vs. CDR accuracy trade-offs during WC-15% validation. Epoch 4 is chosen for the subsequent analysis.
